# Hidradenitis Suppurativa Patients Exhibit a Distinctive and Highly Individualized Skin Virome

**DOI:** 10.1101/2023.10.30.564771

**Authors:** Daan Jansen, Lene Bens, Jeroen Wagemans, Sabrina I. Green, Tom Hillary, Tine Vanhoutvin, An Van Laethem, Séverine Vermeire, João Sabino, Rob Lavigne, Jelle Matthijnssens

## Abstract

Hidradenitis suppurativa (HS) is a chronic inflammatory disease characterized by recurring painful skin lesions. Despite ongoing research, the exact cause underlying the initiation and progression of disease remains unknown. While prior research has linked the skin microbiota to HS pathology, the role of viruses has remained unexplored. To investigate the skin virota, metagenomic sequencing of viral particles was performed on 144 skin samples from 57 individuals (39 HS patients and 18 controls). It was found that the virome is not only linked to BMI, but also to the presence and severity of HS, marking a diverging viral profile in the progression of disease. Despite no differences in alpha-diversity, HS patients exhibited a significantly higher beta-diversity compared to healthy controls, indicating a more personalized virome with reduced viral sharing among patients. We identified distinct groups of commonly shared phages, referred to as the core phageome, associated with either healthy controls or patients. Healthy controls displayed a higher abundance of two core *Caudoviricetes* phages predicted to infect *Corynebacterium* and *Staphylococcus*, comprising normal skin commensals. In contrast, HS patients carried previously uncharacterized phages that were more prevalent in advanced stages of the disease, which likely infect *Peptoniphilus* and *Finegoldia*, known HS-associated pathogens. Interestingly, genes involved in superinfection exclusion and antibiotic resistance could be found in phage genomes of healthy controls and HS patients, respectively. In conclusion, we report the existence of distinct core phages that may have clinical relevance in HS pathology by influencing skin bacteria through mechanisms such as superinfection exclusion and antibiotic resistance.

## INTRODUCTION

Hidradenitis suppurativa (HS) is a chronic inflammatory skin disease of unknown origin that usually develops in skin folds, such as the inguinal and axillary regions [1]. The disease is systemic in nature and is frequently associated with comorbidities, including but not limited to metabolic syndrome, spondylarthritis and inflammatory bowel disease (IBD) [2–4]. The etiology of HS is complex and involves a combination of factors including the environment, lifestyle, hormone imbalances, genetics and the microbiota [5–9]. The interplay between these factors can cause the initiation and development of inflammation around the hair follicles, leading to follicular plugging, abces formation, tissue damage, and ultimately, scar formation [5,10].

Of these factors, the microbiota is of particular interest due to its intricate and enigmatic role in the initiation and development of HS [11]. The human microbiota is known as a complex ecosystem of microorganisms, including viruses, fungi, bacteria, archaea and protozoa, that inhabit the skin and various other mucosal environments. These microorganisms establish a commensal relationship with the human host and provide benefits, such as protecting against invading pathogens. However, this delicate balance is sometimes disrupted, leading to a state called dysbiosis. Dysbiosis has been linked to numerous diseases, including IBD, asthma, depression [12–14], and various skin conditions such as psoriasis, atopic dermatitis and acne vulgaris [15–17].

In this context, a growing body of evidence has linked HS to changes in the bacterial component of the skin microbiota [8,18–22]. These changes are found in various skin regions, including dry, moist and sebaceous areas [23]. HS is a skin condition that commonly affects the skin folds surrounding moist skin regions and is typically characterized by a higher proportion of anaerobic bacteria, such as *Peptoniphilus*, *Porphyromonas*, *Prevotella*, *Finegoldia* and several other microbes [8,19,21,22]. In comparison, healthy individuals tend to have a high proportion of aerobic bacteria that are naturally found in the skin as commensals, including *Corynebacterium, Cutibacterium* and *Staphylococcus* [18,20,21,23]. HS is not linked to a single bacterium, but rather to a mixture of bacteria that are associated with the development and progression of the disease. In that regard, there have been reports of changes in diversity, but these findings are not always consistent [19,21,22]. While some studies indicate no differences [22], others suggest a higher bacterial alpha diversity on the skin of patients seemingly associated with increasing severity of disease [19,21].

While bacterial skin dysbiosis is a frequently described event in the pathophysiology of HS, the skin virota (the collection of viruses infecting human cells, as well as bacteriophages infecting resident bacteria) remains largely unstudied [24]. Nevertheless, the unexplored skin virota could have a vital role, as bacteriophages are capable of controlling bacterial populations through both lytic and lysogenic infection [25–27]. While lytic infection leads to the lysis of bacterial cells and reduces the number of bacteria in a population, lysogenic conversion (the integration of the phage genome in the bacterial genome) allows for the transfer of new genes into the bacterial genome that can influence its metabolism, virulence and antibiotic resistance [28–31]. Furthermore, eukaryotic viruses have the ability to directly infect the human skin, activate the immune system and cause disease, as demonstrated by the association between papillomaviruses, polyomaviruses and herpesviruses, and various dermatological disorders [32–35].

In the present study, we aim to investigate the potential role of the skin virome (both eukaryotic and prokaryotic viruses) in HS patients. To achieve this, we conducted an in-depth analysis of skin swabs (*n=144*) collected from a longitudinal comparative study consisting of 39 patients and 18 healthy controls. By examining these samples, we identified a distinct viral signature that enriches our understanding of the role of the skin microbiome as whole in the pathophysiology of HS.

## METHODS

### Ethical approval

The study was approved by the ethical commission of UZ Leuven (KU Leuven, reference number: S62301). Participants provided a signed informed consent to participate in the study. The design of the study was in accordance with the Declaration of Helsinki and Belgian privacy law.

### Study design and sample collection

The longitudinal comparative study consisted of 18 healthy individuals and 39 hidradenitis suppurativa (HS) patients. Healthy controls were included based on the absence of any dermatological illness and were carefully matched, as closely as possible, for host variables that could potentially impact the microbiota, such as age, sex, ethnicity and others (Supplementary Table 1). Skin samples were collected from both patients (lesions) and healthy controls using ESwabs (1 mL Liquid Amies, Copan, Italy). To ensure a comprehensive dataset, patients were sampled at various locations on the body where skin was affected, at different time points (ideally at baseline, six months and 12 months). A total of 144 samples were collected, consisting of 108 patient samples and 36 healthy control samples. A more detailed overview of the study design and sample collection was previously described by Bens and colleagues [36].

### Viral metagenomics

To perform viral metagenomics on skin swabs, the NetoVIR protocol was used as described before [37]. The skin samples were homogenized using a tabletop vortex and the complete volume (1 mL) was transferred to a 1.5 mL receiver tube. The homogenate was centrifuged at 17,000 g for 3 min, and the resulting supernatant was passed through a 0.8 µm PES filter (Sartorius). A nuclease treatment was then performed using micrococcal nuclease (New England Biolabs) and benzonase (Novagen) at 37°C for 2h. Viral nucleic acids were extracted using the QIAMP® Viral RNA mini kit (60 µL, Qiagen, Venlo, Netherlands) without the addition of carrier RNA. This extraction kit has been previously shown to extract both RNA and DNA viral genomes without introducing any major bias [38]. The Complete Whole Transcriptome Amplification kit (WTA2, Sigma-Aldrich) was employed with minor adaptations (94°C for 2 min, and 17 cycles of 94°C for 30 sec and 70°C for 5 min) to perform reverse transcription and random amplification of the extracted nucleic acids. To address concerns regarding the potential amplification bias towards viruses with a circular single-stranded DNA genome induced by multiple displacement amplification (MDA), a number of points need to be emphasized. First, the WTA2 kit only involved a limited MDA step followed by a classical PCR. Second, the NetoVIR protocol was previously benchmarked on a diverse mock-virome encompassing both linear and circular genomes, and did not reveal any indication of such amplification bias [37]. Next, a purification step was performed on the amplified PCR product using the MSB Spin PCRapace kit (Invitek Molecular), followed by a concentration measurement using the Qubit™ dsDNA HS Assay Kit. Sequencing libraries were prepared from the purified PCR product using the Nextera XT DNA Library kit (Illumina). After library preparation, the sequencing libraries (25 µL) were purified using Agentcourt AMPure XP beads (15 µL, ratio_beads/dsDNA_=0.6, Beckman Coulter) to remove impurities and obtain high-quality libraries. The purified libraries were then assessed for their fragment sizes using the High Sensitivity DNA kit on the Bioanalyzer 2100 (Agilent Technologies). After applying the NetoVIR protocol, a total of 138 high-quality sample libraries were obtained for sequencing. The VIB Nucleomics core performed the sequencing of the libraries using a NovaSeq6000 S1 sequencer (2×150bp, paired end) as the final step. Following the sequencing process, a total of five additional samples were excluded due to a low number of raw reads, indicating low-quality samples, resulting in a total of 133 datasets for further bioinformatic analysis. An elaborate description of the bioinformatic processing, viral identification and classification, diversity analyses and other methodologies can be found in the Extended Data Methods.

## RESULTS

### The skin virome of a comparative hidradenitis suppurativa cohort is characterized by a high proportion of unclassified phages

Skin swabs (*n=144*) were collected from patients (*n=39*) and healthy controls (*n=18*) in a comparative HS cohort (*n=57*). Patients were categorized into three groups based on the severity of the disease, namely Hurley stage 1 (*n=10*), Hurley stage 2 (*n=19*) and Hurley stage 3 (*n=10*). To obtain a complete dataset, patients were sampled at multiple areas of their body where skin was affected (axillary, abdomen, chest, gluteal and groin and genitals), at different time intervals (ideally at baseline, six months and 12 months). Healthy control samples were collected once from the axillary (*n=18*, 50% of samples) and groin and genitals (*n=18*, 50% of samples) to match the vast majority of the patients’ samples (*n=82*, 76.0% of samples; Supplementary Table 1). Of note, 13 of the included patients were diagnosed with IBD, which enabled a comparison of the skin virome of HS patients with and without IBD (Extended Data Methods; Supplementary Table 1).

Next, the virome was characterized by enriching and sequencing viral genomes using the NetoVIR protocol [37]. Bioinformatic analyses on 4.1 billion paired end reads (0.62 TB, 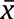=28.6 million reads per sample) were performed to obtain a high-quality (HQ) viral dataset (“HQ Viral NR-Contigs”; Extended Data Figure 1). The HQ viral data set revealed a wide range of recovered viruses across samples, encompassing only 40.2% of the obtained HQ reads (Extended Data Methods; Extended Data Figure 2). This observation is likely attributed to factors such as incomplete public skin phage databases as well as genome fragmentation, corroborating the findings of Graham and colleagues [28]. To include a greater number of partially sequenced phage genome in our analysis and obtain a more representative picture of the skin virome in the complete HS cohort (Healthy+HS), we employed a clustering approach. This method grouped non-redundant contigs into clusters at a “family-like” similarity level using Markov clustering. All clusters including one or more high-quality phages (Extended Data Methods, Extended Data Figure 1) were considered as phage “family-like” viral clusters (FLVC). The FLVC approach increased the retained number of quality-controlled reads to 66.9% (Extended Data Figure 2). Regardless of used viral identification approach (individual HQ-contigs or FLVCs), less than half of bacteriophage contigs could be taxonomically classified at class-level or lower (HQ-contigs_classified_=25.5%, FLVC_classified_=40.2%; Extended Data Figure 3), thereby highlighting the high proportion of previously undescribed phages. Most of the classified skin phage genomes belonged to *Caudoviricetes* (HQ-contigs_classified_=19.3%, FLVC_classified_=39.1%), with only a small fraction classified as *Malgrandaviricetes* (HQ-contigs_classified_=6.2%, FLVC_classified_=1.10%).

### A small and diverse eukaryotic virome with anelloviruses confined to patients

The eukaryotic virome was relatively small, with eukaryotic viruses found in up to 51.2% of skin samples (or 66.6% of individuals), representing 15.9% of the quality-controlled viral reads (FLVC approach; Extended Data Figure 2). In contrast, phages were far more abundant and accounted for 84.1% of the quality-controlled viral reads (FLVC approach; Extended Data Figure 2). The four major eukaryotic viral families (prevalence ≥15%) recovered were *Papillomaviridae*, *Totiviridae*, *Alphaflexiviridae* and *Anelloviridae* (Extended Data Figure 4A). These families consisted of viruses that infect humans (*Papillomaviridae)*, fungi (*Totiviridae*), plants (*Alphaflexiviridae*), as well as small circular viruses with unknown host (*Anelloviridae)*. Despite the frequent occurrence of these viral families, they did not reveal a significantly different relative abundance (RA) between healthy controls and HS patients (*n=38*, Mann-Whitney U, Adj≥P0.05; Extended Data Figure 4B; Supplementary Table 3). A noteworthy observation from our study was that anelloviruses, for which the host cells and biological function in the human body are still unknown [39], were exclusively detected in skin samples of HS patients. In addition, the detected totiviruses were predicted to infect the fungi *Malassezia*, which typically exist as a commensal on human skin surfaces [40]. Another peculiar finding included the detection of members of the *Dicistroviridae* and *Trichomonasvirus* capable of infecting the house dust mite and the protozoan parasite *Trichomonas*, respectively, in several of the patients. Ultimately, our findings suggest that eukaryotic viruses are unlikely to play a significant role in HS pathology due to their low abundance and inability to differentiate between healthy and diseased individuals.

### The skin phageome composition is associated with the presence and severity of hidradenitis suppurativa

To investigate the potential link between HS pathology and the phageome, we analyzed the compositional variation of the phageome in the HS cohort. In doing so, a stepwise distance-based redundancy analysis (dbRDA) was used to identify factors associated with the phageome composition of the complete HS cohort (Healthy+HS; Supplementary Table 4). First, the composition was found to be significantly associated with both categorical BMI (cBMI) and disease status (i.e., healthy or HS patients), thus confirming the link between HS pathology and skin phages *(n=51*, multivariate dbRDA, FLVC-level, cBMI R_2_=5.16%, disease status R_2_=1.63%, AdjP<0.05; Figure 1a,b). The fact that healthy controls were not fully matched, and an association between cBMI and disease status was present (*n=51*, χ^2^=15.0, r=0.736, AdjP=0.00182; Figure 1c; Supplementary Table 5), suggest that cBMI may act as a confounding variable on further analyses. Second, it was observed that in the univariate analyses, age and geographical location (i.e., province) also had a small influence on the skin phageome, even though their explanatory power was weaker compared to both cBMI and the presence of skin disease (*n=51*, univariate dbRDA, FLVC-level, cBMI R_2_=5.16%, disease status R_2_=3.32%, province R_2_=2.69%, age R_2_=1.56%, AdjP<0.05; Figure 1b). Third, as mentioned earlier, we aimed to assess the impact of a concurrent IBD diagnosis on the phageome. Nevertheless, this analysis did not reveal any discernible differences in the skin phageome between patients with IBD and those without.

**Figure 1:**
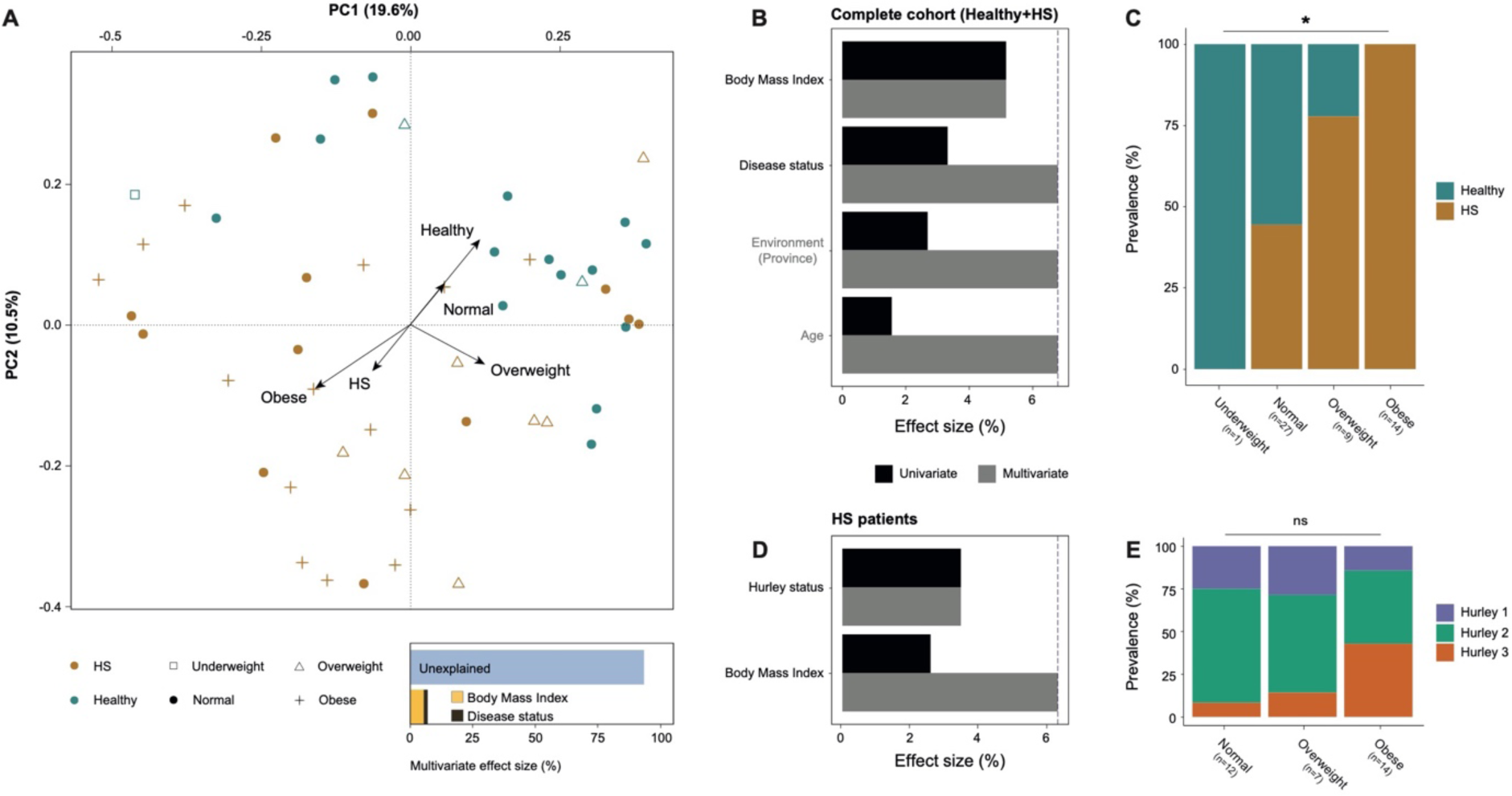
Skin virome variation in the complete and patient study cohort. **A,** A Principal Coordinate Analysis visualizing the inter-individual differences of the skin virome composition (*n=51*, FLVC level, Bray-Curtis dissimilarity) of the complete cohort (Healthy+HS), coloured by disease status and with shapes depicted by categorical BMI. The barplot at the bottom illustrates the significant covariates of the skin virome (93.2% unexplained); arrows on the plot indicate the effect sizes of these significant covariates (underweight (*n=1*) not shown). **B, C,** Panels show the metadata variables significantly associated with the skin virome composition in the complete cohort (*n=51*, HS+Healthy) and in HS patients (*n=33*) only, respectively (dbRDA, FLVC level, Bray-Curtis dissimilarity). The effect sizes of covariates are shown independently as univariate analysis (in black) or as a cumulative model (in grey). **D,E,** Panels illustrate the associations between disease status (*n=51,* chi-squared test, AdjP<0.05) or disease severity (*n=33*, chi-squared test, AdjP≥0.05) and categorical BMI. Multiple testing adjustment (Benjamini-Hochberg method) was performed, and significant associations (AdjP<0.05) are represented by an asterisk (*). Abbreviations: Hidradenititis suppurativa (HS)/Healthy controls (Healthy), complete cohort (Healthy+HS) and non-significant (ns).

Our next goal was to investigate the potential link between the disease severity (i.e., Hurley stages) and the composition of the skin phageome. To achieve this, the previous dbRDA analysis was restricted to the samples pooled per HS patient (excluding healthy controls) (Figure 1d). As expected, the phageome composition remained significantly associated to both disease severity and cBMI *(n=33*, multivariate dbRDA, FLVC-level, disease severity R_2_=3.44%, cBMI R_2_=2.81%, AdjP<0.05; Figure 1d Supplementary Table 6). It is important to note that in this particular analysis, no significant association was found between disease severity and cBMI, indicating that cBMI does not serve as a confounding variable in this subgroup (*n=33*, χ^2^=4.80, r=0.520, AdjP=0.308; Figure 1e; Supplementary Table 5). Taken together, these findings suggest that together with cBMI, disease presence and severity were key covariates of the phageome composition, marking a diverging viral profile in the progression of HS pathology.

To obtain insights in the longitudinal and spatial differences of the phageome composition, the previous analysis was repeated on the complete HS cohort, at the individual sample level (Supplementary Table 7). The phageome was found to be significantly linked with cBMI and disease status (*n=99*, multivariate dbRDA, FLVC-level, cBMI R_2_=1.06%, disease status R_2_=0.650%, AdjP<0.05), consistent with previous findings. While no longitudinal effect was observed, skin location was identified as a non-redundant explanatory variable with minor effect size (*n=99*, multivariate dbRDA, FLVC-level, R_2_=1.06%, AdjP=0.004). Despite our expectations, most therapies and types of skin lesions did not reveal any relationship with the phageome composition, except for metformin treatment and skin abscesses, which only showed a minimal association (*n=99*, multivariate dbRDA, FLVC-level, abscess R_2_=0.640%, metformin R_2_=0.420%, AdjP<0.05). These analyses emphasize the importance of considering major confounding variables (effect size > 1%), such as cBMI, wherever appropriate in subsequent analysis to accurately establish the precise dynamics between the phageome and HS pathology.

### Hidradenitis suppurativa patients exhibit a distinctive skin virome that is highly individualistic and is marked by a reduced healthy core phageome

Having established the link between the phageome and HS pathology, our focus shifted towards examining how skin phages might differentiate healthy controls from disease. For this purpose, a BMI-deconfounded cohort (only using patients and controls with cBMI=normal) was set up, consisting of 12 HS patients and 15 healthy controls (Figure 2A). Subsequent diversity analyses did not show a significant difference in alpha-diversity (i.e., Shannon diversity) between HS patients and healthy controls (*n=27*, Mann-Whitney U, FLVC-level, r=0.197, AdjP=0.323; Figure 2B; Supplementary Table 8). However, HS patients exhibited a significantly higher beta-diversity (i.e. Bray-Curtis dissimilarity) compared to healthy controls (*n=27*, Mann-Whitney U, FLVC-level, r=0.552, AdjP=1.00e-12; Figure 2C; Supplementary Table 8). This meant that patients had a far more personalised skin phageome in comparison to the healthy controls. To corroborate this finding, we determined the degree of individuality by measuring the proportion of phage FLVC that were unique to a single individual (referred to as ‘unique FLVC’). Healthy controls had a total of 17.4% unique FLVC, while patients had a total of 46.5% unique FLVC (Figure 2D,E). In contrast, 82.6% of FLVC were shared among healthy controls, whereas only 53.5% were shared among HS patients. To explore viral sharing in greater detail, we focused on phage FLVC that were detected in 50% or more of either healthy individuals or HS patients (Figure 2D,E). These FLVCs will be referred to as the “core skin phageome” further on. A total of 13 core FLVCs (cFLVC) were identified in the BMI-deconfounded cohort (Supplementary Table 9). Among them, four cFLVC were found in both healthy controls and HS patients, while 7 cFLVC and two cFLVC were exclusively found in healthy controls and HS patients, respectively (Supplementary Table 9). In addition, a similar analysis was conducted to investigate cFLVC among different Hurley stages, which showed that cFLVC4 and cFLVC5 were present in all of the patients with severely progressed skin disease (Hurley stage 3; Supplementary Table 10). Finally, alpha and beta-diversity analyses were also conducted on the non-BMI deconfounded cohort, revealing similar outcomes, underscoring a minimal significance of BMI in the results (Extended Data Figure 5; Supplementary Table 8,9).

**Figure 2:**
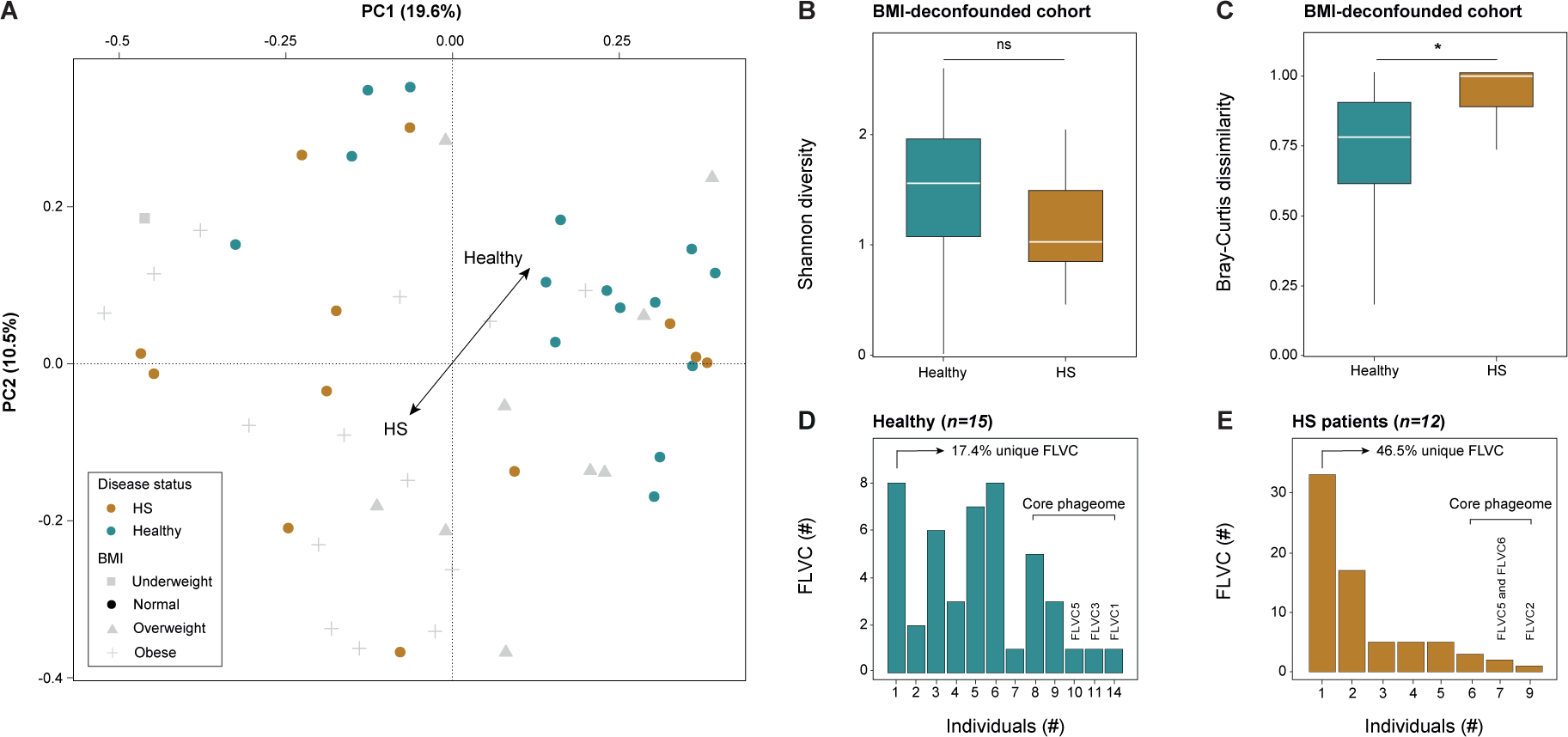
Skin virome diversity and viral sharing in the BMI-deconfounded study cohort. **A,** A principal Coordinate Analysis visualizing the inter-individual differences of the skin virome composition (*n=51*, FLVC level, Bray-Curtis dissimilarity) of the complete cohort (Healthy+HS) coloured by disease status and with shapes depicted by categorical BMI. Individuals whose BMI fall outside the normal range (17.5-25 kg/m^2^) are shaded in grey. **B,** Alpha diversity (Shannon index) boxplot stratified according to disease status in BMI-deconfounded cohort (*n=27*, FLVC level, BMI=normal, Mann-Whitney U, AdjP≥0.05). **C,** Beta diversity (Bray-Curtis dissimilarity) boxplot stratified according to disease status in BMI-deconfounded cohort (*n=27*, FLVC level, BMI=normal, Mann-Whitney U, AdjP=1.00e-12). **D,** Barplot showing the absolute and relative number of FLVCs shared within healthy individuals in BMI-deconfounded cohort (*n=15,* BMI=normal). **E,** Barplot showing the absolute and relative number of FLVCs shared between HS patients in BMI-deconfounded cohort (*n=12,* BMI=normal). The core virome is defined by FLVCs shared between 50% or more of healthy individuals or HS patients (three most shared viruses are listed). Multiple testing adjustment (Benjamini-Hochberg method) was performed, and significant associations (AdjP<0.05) are represented by an asterisk (*). Abbreviations: Hidradenititis suppurativa (HS)/Healthy controls (Healthy), complete cohort (Healthy+HS), family-like viral cluster (FLVC) and non-significant (ns).

Taking these findings into account, we hypothesized that the phage cFLVC could distinguish the skin of healthy controls from that of HS patients, and subsequently compared the RA of the 13 previously identified cFLVC between both groups. Healthy controls showed a significantly higher RA of both cFLVC1 and cFLVC3 compared to HS patients (*n=51*, Mann-Whitney U, FLVC-level, AdjP<0.05; Figure 3a; Supplementary Table 11). In comparison, none of the cFLVC were found to be significantly higher among HS patients compared to healthy controls (*n=51*, Mann-Whitney U, FLVC-level, AdjP≥0.05; Figure 3a; Supplementary Table 11). Upon classification of these cFLVC, it was observed that cFLVC1 and cFLVC3 bore no resemblance to phages in RefSeq (Extended Data Figure 6). However, cFLVC3 contained a few members showing similarities as high as 95.7% across 50% of their genomes to “uncultured *Caudovirales* phage” genomes in public databases (Supplementary Table 12), suggesting that they represent a group of poorly described phages within the *Caudoviricetes* class with lysogenic potential. Collectively, these results suggest the existence of a core phageome in healthy individuals, which is depleted in patients with HS.

**Figure 3:**
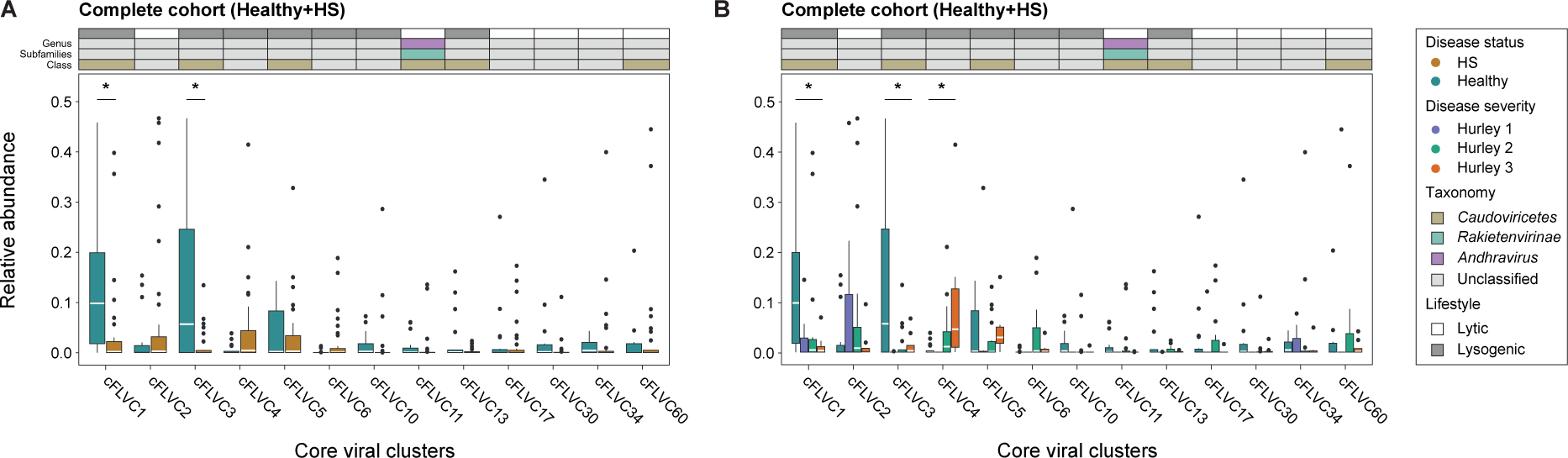
The relative abundance of the cFLVCs in the complete study cohort. **A,** Relative abundance boxplot of the cFLVCs stratified according to disease status in the complete cohort (*n=51*, FLVC level, Mann-Whitney U, AdjP<0.05). The core virome is defined by FLVCs shared between 50% or more of healthy individuals or HS patients. **B,** Relative abundance boxplot of the cFLVCs stratified according to disease severity in the complete cohort (*n=51*, FLVC level, Kruskal-Wallis, AdjP<0.05). Classification (AS≥0.1) and lifestyle of representative high-quality phages of cFLVCs are shown (top) at class, subfamily and genus taxon. Multiple testing adjustment (Benjamini-Hochberg method) was performed and significant associations (AdjP<0.05) are represented by an asterisk (*). Abbreviations: Alignment score (AS), Hidradenititis suppurativa (HS)/Healthy controls (Healthy), complete cohort (Healthy+HS), family like viral cluster (FLVC) and non-significant (ns).

### Patients with advanced disease are associated with an unidentified phage cluster

Following the failure to identify a cFLVC positively associated with HS, it was suggested that a high degree of patient individuality, coupled with the influence of disease progression, could obscure significant associations. For that reason, the RA of the 13 previously identified cFLVC was compared between various stages of disease severity (Figure 3b; Supplementary Table 13, 14). Despite the lack of significant differences in RA between the various disease stages for cFLVC1 and cFLVC3 (Kruskal-Wallis test, FLVC-level, AdjP≥0.05), a notably lower RA was identified when compared to healthy controls, consisting with previous findings (Kruskal-Wallis with post-hoc Dunn test, FLVC-level, AdjP<0.05). A significant association between disease severity and cFLVC4 was observed, displaying a gradual rise in RA with the progression of the disease (*n=51*, Kruskal-Wallis, FLVC-level, r=0.334, AdjP=0.00408; Fig. 3B; Supplementary Table 13). More specifically, the RA of cFLVC4 was significantly higher in advanced disease (i.e., Hurley stage 3, *n=26*, post-hoc Dunn test, AdjP=0.00591), but not in mild and moderate disease (i.e., Hurley stage 1 and 2, post-hoc Dunn test, AdjP≥0.05), compared to healthy controls. Furthermore, patients with moderate and advanced disease showed a significantly higher RA of cFLVC4 compared to mild disease (post-hoc Dunn test, AdjP<0.05), but were not statistically different from each other (post-hoc Dunn test, AdjP≥0.05). A more detailed analysis of phage genomes in cFLVC4 showed no similarities to phage genomes in RefSeq (Extended Data Figure 6) or other public databases (Supplementary Table 12), suggesting that it corresponds to a previously unidentified phage cluster with the capability of lysogenic conversion (Supplementary Table 12). Taken together, these results point toward the existence of a core phage FLVC that gains more ground in the later stages of HS.

Finally, the aforementioned analysis was replicated using only a single sample per patient (baseline sample) in a disease location-deconfounded sample set (originating from the axillary and groin and genital region). In this cohort, sample location was not associated to disease severity and therefore did not act as a confounding variable (*n=55*, Chi-squared test, AdjP=0.985; Extended Data Figure 7a; Supplementary Table 15). The RA of all three cFLVC (cFLVC1, cFLVC3 and cFLVC4) were still found to be associated with the severity of HS (*n=55*, Kruskal-Wallis test, FLVC-level, AdjP<0.05; Extended Data Figure 7b,c; Supplementary Table 16). Interestingly, the results could not be reproduced when focussing solely on either the axillary or groin and genital region, likely due to the reduced statistical power from small samples sizes (Kruskal-Wallis test, FLVC-level, AdjP≥0.05; Extended Data Figure 8; Supplementary Table 16).

### The healthy core phageome is predicted to infect *Corynebacterium* and *Staphylococcus* and is marked by a high presence of prophage-mediated bacterial defence systems

Having identified the phages associated with a healthy skin, our next goal was to investigate how these phages might influence the skin microbiome. To do so, we used an *in silico* host prediction tool (i.e., RaFAH) to forecast the likely bacterial host. Based on this, it was predicted that cFLVC1 phages target Actinobacteria, whereas cFLVC3 phages infect Firmicutes (Figure 4A; Supplementary Table 12). To obtain a more refined taxonomical resolution, *in silico* host predictions were repeated to estimate the bacterial genus. The findings indicated that cFLVC1 phages primarily target *Corynebacterium* species, while cFLVC3 phages predominantly infect *Staphylococcus* species (Figure 4A; Supplementary Table 12). Furthermore, 16S bacterial data obtained from a parallel study was used to confirm the association between the healthy core phages and their respective bacterial host (Figure 4B; Supplementary Table 17) [36]. A notable positive correlation was found between the RA of cFLVC1 and the RA of *Corynebacterium* (*n=51*, r=0.29, AdjP=0.039; Figure 4B; Supplementary Table 17), providing further evidence of a connection between these phages and their bacterial hosts. Moreover, no significant correlation was detected between the RA of cFLVC3 and the RA of *Staphylococcus* (*n=51*, AdjP=0.052; Figure 4B; Supplementary Table 17). This lack of a significant correlation may be due to the diverse nature of *Staphylococcus* on the skin [41], which include both symbiotic (i.e., *Staphylococcus epidermidis*) and pathogenic species (i.e., *Staphylococcus aureus)*, that could be associated with phages other than cFLVC3. In addition, the bacterial genus *Escherichia*-*Shigella*, which served as a negative control for the correlation analysis, was found to have no significant association with either cFLVC1 or cFLVC3 (*n=51*, AdjP<0.05; Figure 4B; Supplementary Table 17).

**Figure 4:**
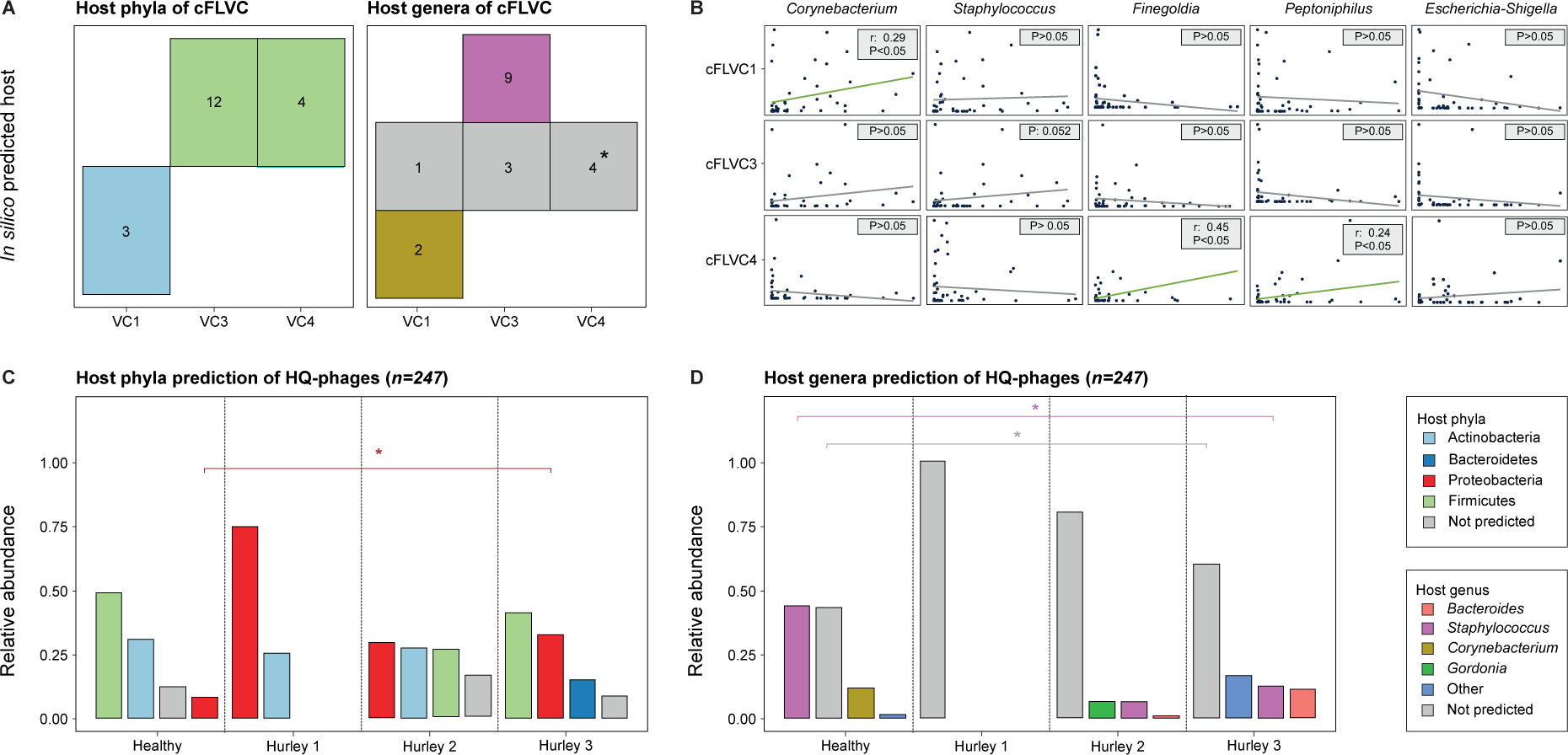
Host prediction of the skin virome. **A,** Tile plot of *in silico* predicted bacterial host phyla (left) and genera (right) of high-quality phage representative sequences of the core viral clusters significantly associated with disease status and disease severity. The number of representatives of each FLVC are depicted in each tile and are near-complete (>50% complete) phage sequences. The asterisk denotes that a comparison of phage-incorporated bacterial proteins of the representatives of FLVC4 to protein databases could provide insights into the potential host (i.e., *Finegoldia* and *Peptoniphilus*; Supplementary Table 12). **B,** Comparison of the relative abundance of core viruses to the relative abundance of predicted bacterial genera, as determined by 16S bacterial data. The Escherichia-Shigella group (most abundant genera) was not predicted by former analysis and served as a negative control. **C,** Compositional barplot of *in silico* predicted bacterial host phyla of high-quality phages stratified according to disease severity (*n=43*, Kruskal-Wallis, AdjP<0.05). **D,** Compositional barplot of *in silico* predicted bacterial host genera of high-quality phages stratified according to disease severity (*n=43*, Kruskal-Wallis, AdjP<0.05). *In silico* host predictions are determined with Random Forest Assignment of Hosts (RaFAH). Multiple testing adjustment (Benjamini-Hochberg method) was performed. Abbreviations: family like viral cluster (FLVC).

To complete our analysis of the healthy core phageome, the representative phage genomes of cFLVC1 and cFLVC3 were functionally annotated, as shown in Figure 5A. Notably, these genome maps revealed the predicted presence of proteins that enhance fitness of the bacterial host and were found within the majority of individual phage genomes (i.e., cFLVC1=66.7%, cFLVC3=58.3%; Figure 5B; Supplementary Table 12). These predicted proteins including lipoproteins such as “host cell surface-exposed lipoprotein” and “Siphovirus gp157 protein”, located within the lysogeny module of the phage genomes. These lipoproteins were previously recognized for their involvement in ‘superinfection exclusion’, a mechanism that safeguards the bacterial cell by preventing secondary viral infections [42]. Collectively, the results suggest that phages present on the skin of healthy individuals could integrate their genetic material into the genomes of *Corynebacterium* and *Staphylococcus*, leading to an enhanced fitness of the bacterial host through the mechanisms of superinfection exclusion.

**Figure 5:**
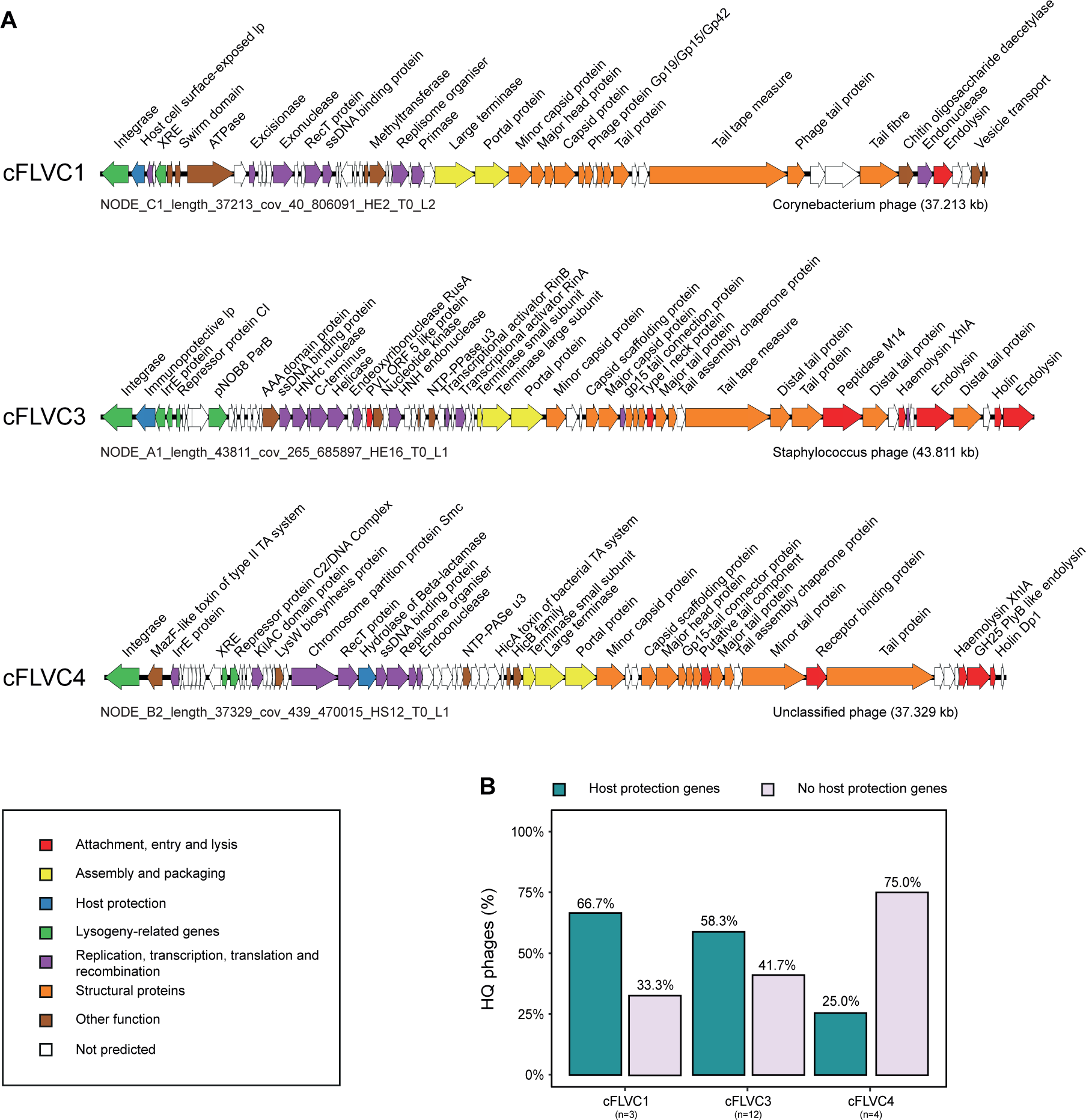
Genome maps of representative high-quality phages for each cFLVC. **A,** Genome maps of representative high-quality (near-complete) phages of FLVC1, FLVC3 and FLVC4 as obtained by Cenote-Taker2 [45] with arrows representing predicted ORFs. The genomes of FLVC1 (NODE_C1_length_37213_cov_40_806091_HE2_T0_L2) and FLVC3 (NODE_A1_length_43811_cov_265_685897_HE16_T0_L1) were determined to be lysogenic *Corynebacterium* and *Staphylococcus* phages, respectively. The representative of FLVC4 (NODE_B2_length_37329_cov_439_470015_HS12_T0_L1) is determined to be a lysogenic phage with unpredicted host genus using RaFAH [46]. Phage genomes are linearized to improve visualization. **B,** Percentage of high-quality phages containing genes advantageous to the bacterial host (by increasing host fitness) stratified according to each cFLVC. Abbreviations: family like Viral cluster (FLVC) and high-quality phages (HQ phages).

Lastly, we conducted an *in silico* host prediction on the complete phageome, expanding our analysis beyond individual viral clusters (Figure 4C,D). The results revealed that, at the phylum taxonomy level, there is a significantly higher RA of proteobacteria-infecting phages in HS patients compared to healthy controls (Kruskal-Wallis with post-hoc Dunn test, AdjP<0.05, Figure 4C; Supplementary Table 18). Analysis of the genus level not only indicated a lower RA of *Staphylococcus*-infecting phages, but also a higher RA of phages without an assigned host in HS patients compared to healthy controls (Kruskal-Wallis with post-hoc Dunn test, AdjP<0.05; Figure 4D; Supplementary Table 18).

### Disease-associated core phages potentially infect *Finegoldia* and *Peptoniphilus* and revealed the presence of antibiotic-resistance genes

As mentioned previously, disease-associated core phages (i.e., cFLVC4) gradually gained more ground in later stages of HS, eventually establishing a presence in all patients with advanced disease. To further understand the impact of these phages on the skin microbiome of patients, we conducted *in silico* host prediction analyses. They revealed that cFLVC4 phages preferred to infect Firmicutes, while accurate predictions regarding the bacterial genus could not be determined (Figure 4A; Supplementary Table 12). As an alternative strategy to gain insights into their potential host, phage-incorporated bacterial proteins found within the phage genomes were explored, such as Haemolysin XhlA or Chromosome partition protein Smc (Supplementary Table 12). These hypothesis was that these proteins, acquired from past infections, could provide valuable insights about the present host. Comparison of such proteins with bacterial databases suggested that cFLVC4 phages likely target *Peptoniphilus* (best blastp hit: AAI%= 99.0%; coverage=96.3%) and *Finegoldia* (best blastp hit: AAI%= 95.3%; coverage=100%), both known pathogens associated with HS [43]. In addition, the RA of both bacterial genera showed a positive and significant correlation with the RA of cFLVC4 phages (*n=51*, r>0.30, AdjP<0.05; Figure 4B; Supplementary Table 17), adding further evidence to support the relationship between these phages and their host. Lastly, after carefully analysing the genome maps of the disease-associated core phages, we observed the presence of antibiotic-resistance genes, although only a minority of representative phage genomes (i.e., Hydrolase of Beta-lactamase; cFLVC4=25.0%, Figure 5B; Supplementary Table 12). These genes enable the bacterial host to improve its fitness by acquiring resistance against beta-lactam antibiotics. Taken together, the collective evidence suggests that phages present in advanced disease may integrate their genetic material into the genome of *Peptoniphilus* and *Finegoldia*. This integration process might result in enhanced fitness of the bacterial host by providing them with antibiotic resistance genes, which emphasizes the potential clinical relevance of phages in the pathology of HS.

## DISCUSSION

To the best of our knowledge, this research represents the first investigation of the skin virome in patients diagnosed with hidradenitis suppurativa. Here, we found a small yet diverse eukaryotic virome with anelloviruses confined to disease. Earlier studies have already indicated that anelloviruses were rarely found on the skin of healthy individuals [44], suggesting a logical relationship with the inflammatory component (i.e., immune cells) present in patients but not healthy individuals. Despite this, it remains unlikely that eukaryotic viruses play an important role in HS pathology, given their low abundance and inability to differentiate between healthy and diseased individuals. By contrast, when we shifted our attention to the phageome, we found a significant association with HS pathology. This association emerged after employing a viral clustering approach, which allowed us to obtain a more comprehensive representation of the skin phageome. While a majority of the phages could be recognized, only a minority could be classified, indicating the presence of previously uncharacterized phages in our dataset.

When considering the relationship between the phageome and HS pathology, it was evident that the phageome exhibited a robust association with both the presence and severity of HS. This indicated that the skin phageome may possess the ability to differentiate, to a certain extent, between healthy individuals and HS patients. Interestingly, the results revealed that this differentiation was not related to the presence of a concurrent diagnosis of IBD. To further explore the mechanism by which the phageome achieved this differentiation, we examined various diversity indices. This indicated that patients had a significantly more personalized phageome compared to controls, indicating a reduced level of phage sharing among patients. To gain a deeper understanding of phage sharing, the concept of “core viral clusters” was introduced, which comprised phage clusters that were present in 50% or more of the individuals.

Using this concept, two core phage clusters, namely cFLVC1 and cFLVC3, both belonging to the *Caudoviricetes* class, were found to be highly abundant on the skin of healthy individuals but not of HS patients. These phages were identified as lysogenic and capable of infecting *Corynebacterium* and *Staphylococcus*, which comprised known skin commensals. Furthermore, they harboured genes involved in superinfection exclusion, a mechanism that confers bacterial resistance to a secondary infection of closely related phages. Thus, it was hypothesized that cFLVC1 and cFLVC3 may enhance the fitness of *Corynebacterium* and *Staphylococcus*, thereby stabilizing the microbiota and promoting skin symbiosis. Interestingly, other studies have suggested that genes for superinfection exclusion may be constitutively transcribed during the prophage status, highlighting the intricate role of the phageome within the skin microbiome [42]. To conclude our study, we examined different stages of the disease and identified a previously unidentified core phage cluster, namely cFLVC4, that were more prevalent in advanced stages of the disease. This core phage cluster, which was discovered in all patients with late-stage disease (i.e., Hurley stage 3), exhibited a lysogenic nature and likely has the ability to infect *Peptoniphilus* and *Finegoldia*, two known pathogens associated with HS [43]. Remarkably, the phages revealed the presence of antibiotic resistance genes, potentially complicating patient treatment and promoting skin dysbiosis. We speculate that this observation is linked to the higher use of antibiotics in late stage (80%) compared to mild or moderate disease (40% and 31.2%; Supplementary Table 1).

To summarize, our study suggests a relationship between the skin virome and HS pathology. We identified distinct groups of phages associated with either healthy controls or HS patients, which could promote bacterial fitness. In healthy individuals, these phages could contribute to the stability of the skin microbiota and foster symbiosis, through mechanisms like superinfection exclusion. However, in HS patients, the presence of phages could potentially contribute to skin dysbiosis by providing antibiotic resistance genes and complicating treatment, underscoring the clinical significance of the virome in HS pathology.

## DECLARATIONS

### Availability of data and materials

Supplementary Table 1 contains the clinical metadata. The raw sequence data have been deposited to the NCBI Sequence Read Archive with BioProject accession number PRJNA961962. The high-quality phage sequences (representatives of cFLVC1, cFLVC3 and cFLVC4) were deposited to GenBank under the following accession numbers: OQ890309-OQ890326. The ViPER (Virome Paired-End Reads) pipeline was used for bioinformatic processing of the raw paired end reads and is publicly available at GitHub (https://github.com/Matthijnssenslab/ViPER). All the data to reproduce virome analysis are available at https://github.com/Matthijnssenslab/IBDVirome/tree/main/IBDHS.

## Author Contributions

The study was conceived by JM, RL and SV. Experiments were designed by DJ, JW and LB. Sampling was set up by TV and TH. Experiments were performed by DJ, AVL and JS. Bioinformatic and statistical analysis of the sequence reads was performed by DJ. SG, JW, LB, RL, JM and DJ drafted the manuscript. All authors revised the article and approved the final version for publication.

## Acknowledgements

We would like to express our gratitude to Dr. Şimşek and Mr. Chiu for extensive proofreading of the manuscript.

## EXTENDED DATA FIGURES

**Extended Figure 1:**
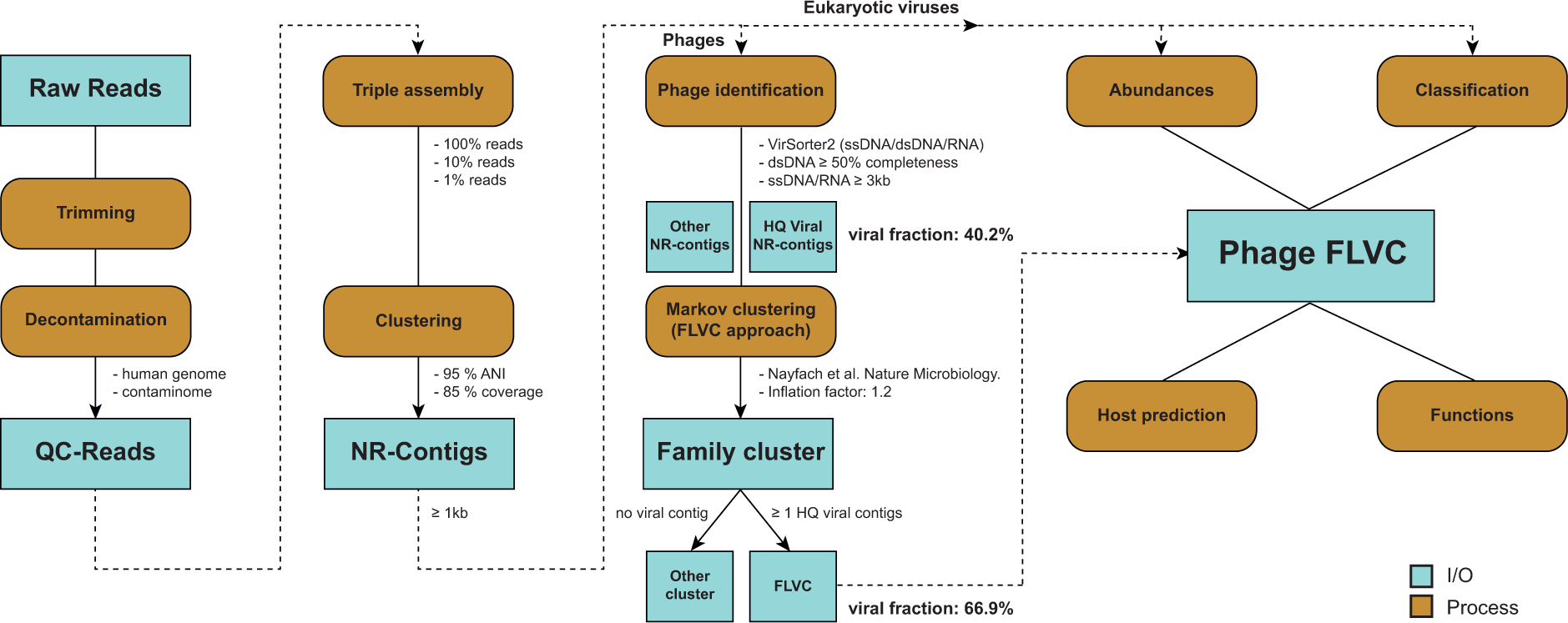
Bioinformatic analysis of the skin virome. An overview of the bioinformatics (dry lab) analysis, also known as drylab analysis, of the skin virome. First, raw reads undergo trimming and decontamination to acquire high-quality reads, referred to as “QC-Reads”. QC-reads are then *de novo* assembled three times, using subsets of reads, resulting in long contiguous sequences through a process called “triple assembly”. Next, contigs are clustered to obtain a non-redundant contig set. Viruses are identified using specific methods outlined in the methods sections, classified and abundances are calculated. Additionally, the functional characteristics of the identified viruses and their hosts are determined through further analysis. Abbreviations: Input/Output (I/O), Contiguous sequences (contigs), non-redundant (NR) and viral cluster (FLVC).

**Extended Figure 2:**
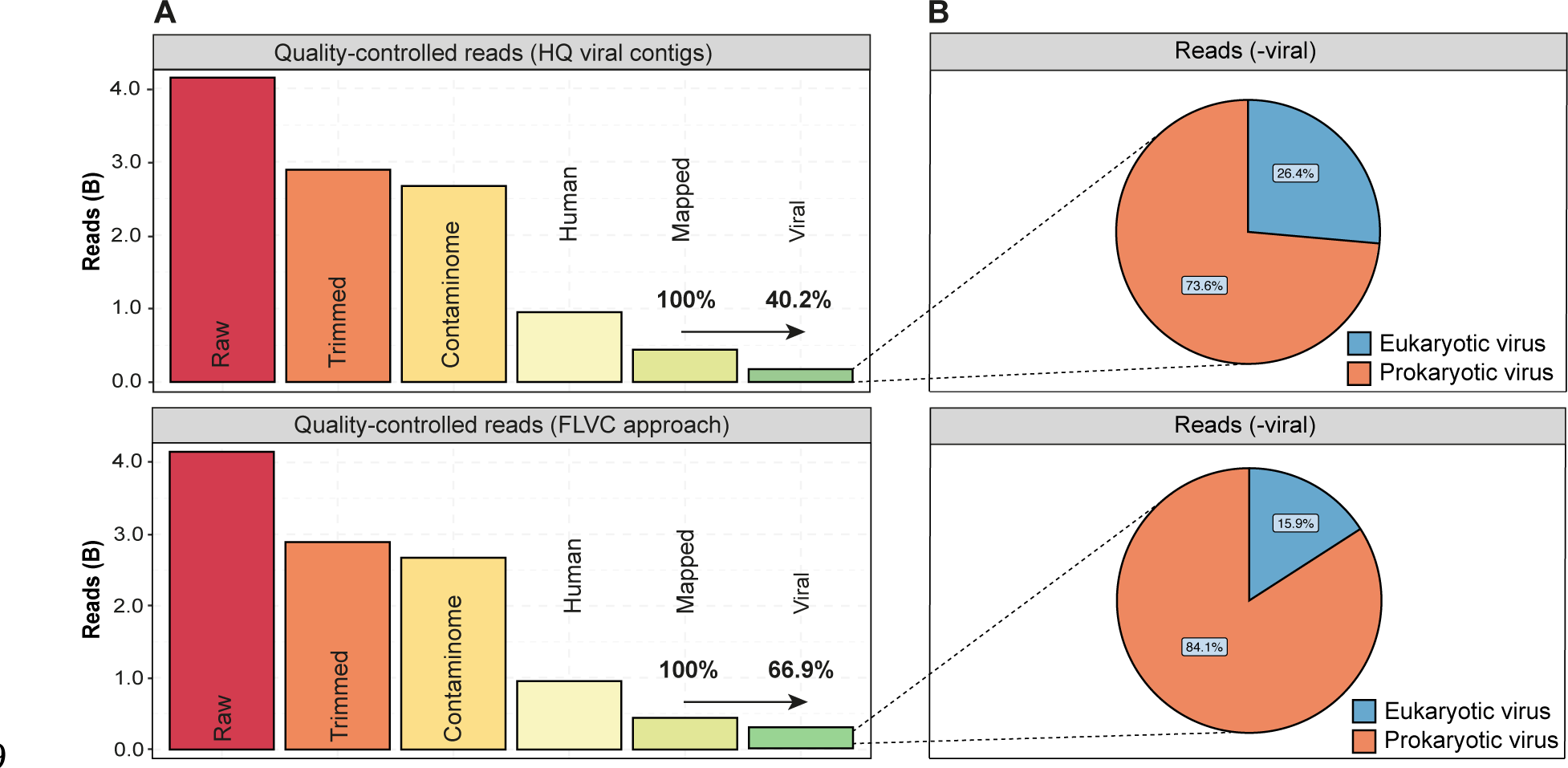
The quality control steps implemented to obtain the viral metagenome of the complete study cohort. **A,** The number of reads obtained during each quality control step for the HS dataset is presented, along with a comparison of identifying only high-quality phages versus using the viral cluster approach to identify phages (raw reads=4.15 billion). Viral reads (4.27%-7.08% of raw reads) represent quality-controlled reads mapped to viruses identified with (number of viral reads=294 million, %viral reads=66.9%) or without (number of viral reads=177 million, %viral reads=40.2%) the viral cluster approach. **B,** Pie chart depicting the distribution of viral reads, mapping to either prokaryotic or eukaryotic viruses, stratified based on the viral identification approaches, as described before. Abbreviations: Hidradenititis suppurativa (HS), Complete cohort (Healthy+HS) and family like viral cluster (FLVC).

**Extended Figure 3:**
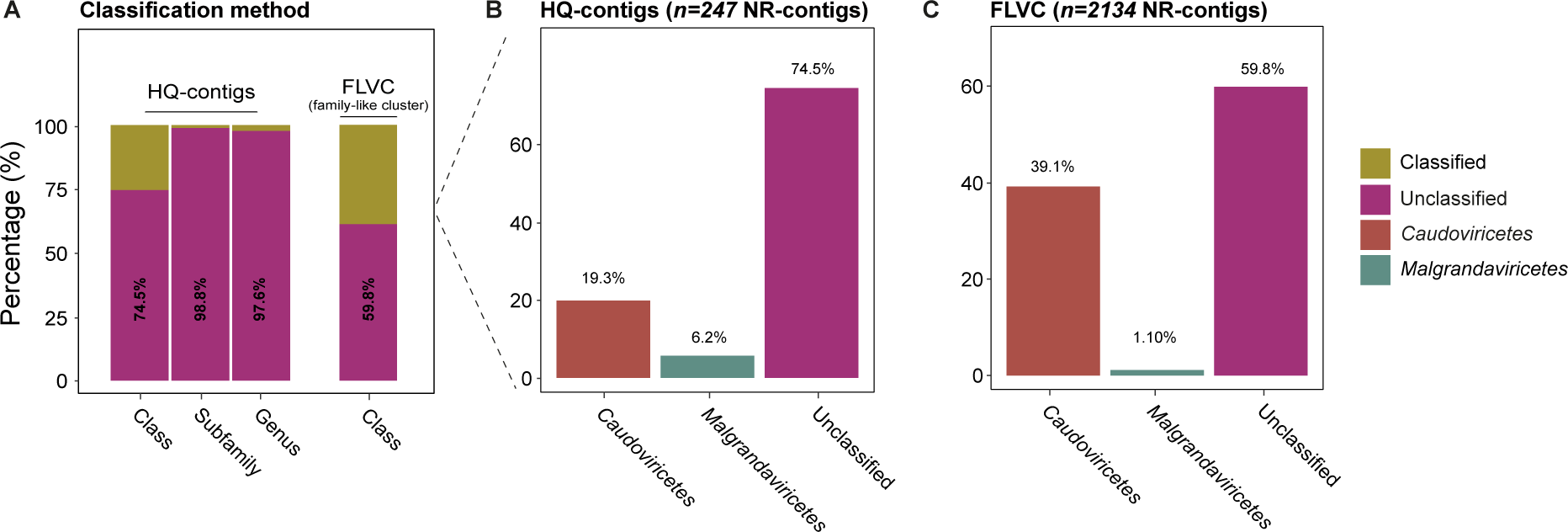
The classified and unclassified phageome in the complete study cohort. **A,** The percentage of classified and unclassified non-redundant phage contigs in the high-quality phage (*n=247* NR-contigs) and FLVCs (*n=2134* NR-contigs) dataset. **B,** The percentage of classified and unclassified non-redundant phage contigs in the high-quality phage dataset on class taxon (25.5% classified and 74.5% unclassified). **C,** The percentage of classified and unclassified non-redundant phage contigs in the viral cluster phage datasets on class taxon (41.2% classified and 59.8% unclassified). The classification of phages was performed by homology-based (AS ≥ 0.1) and marker gene classification (see Methods). Abbreviations: Alignment score (AS), Complete cohort (Healthy+HS), high-quality contigs (HQ-contigs) and non-redundant contigs (NR-contigs).

**Extended Figure 4:**
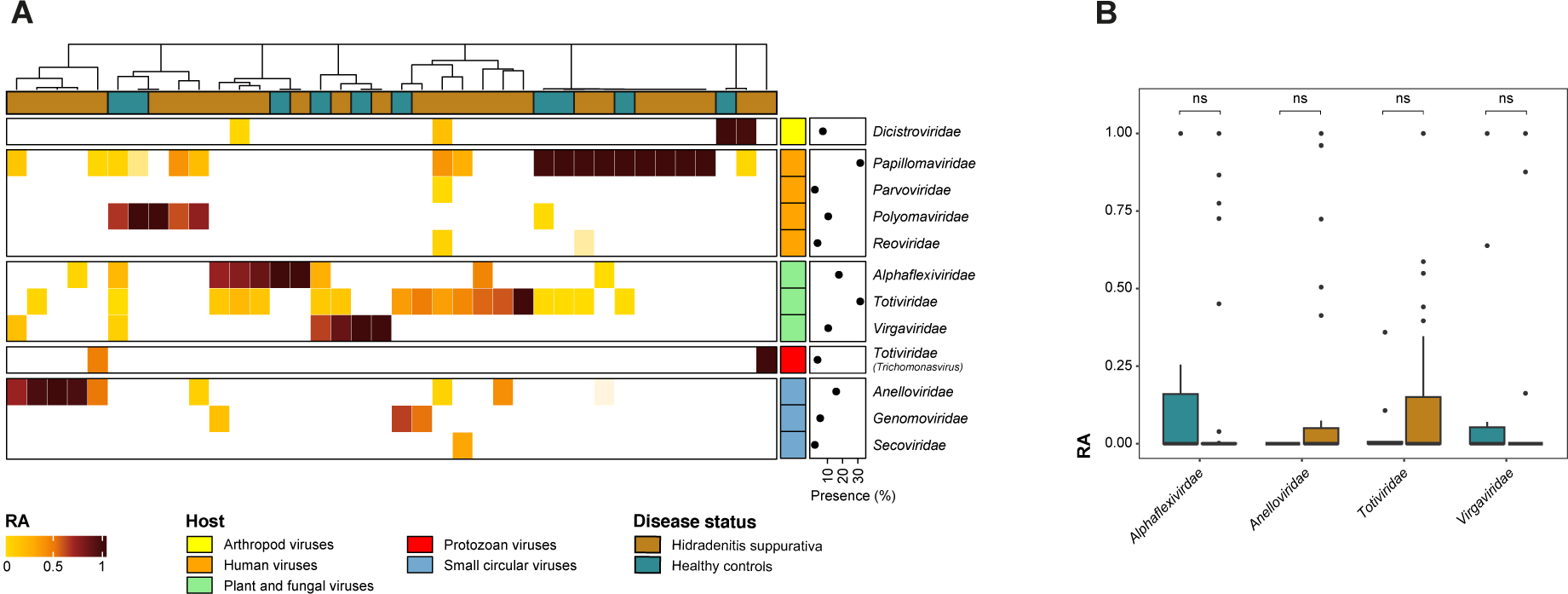
The eukaryotic viral composition in the complete study cohort. **A,** A relative abundance heatmap depicting the eukaryotic virome in HS patients and healthy controls (family level, *n=38*, eukaryotic viral presence=66.7%). The eukaryotic viral families have been categorized into groups based on their hosts, which are either known (plant/fungal and animal viruses) or unknown (small circular viruses). The classification of eukaryotic viruses was performed by homology-based classification (AS ≥ 0.1). **B,** Relative abundance boxplots of the most prevalent viral families (*Alphaflexiviridae*, *Anelloviridae*, *Totiviridae* and *Virgaviridae*) stratified according to disease status in complete cohort (Mann-Whitney U, AdjP≥0.05). Adjustment for multiple testing was performed using the Benjamini-Hochberg method. Abbreviations: Alignment score (AS), Complete cohort (Healthy+HS), Relative abundance (RA) and not-significant (ns).

**Extended Figure 5:**
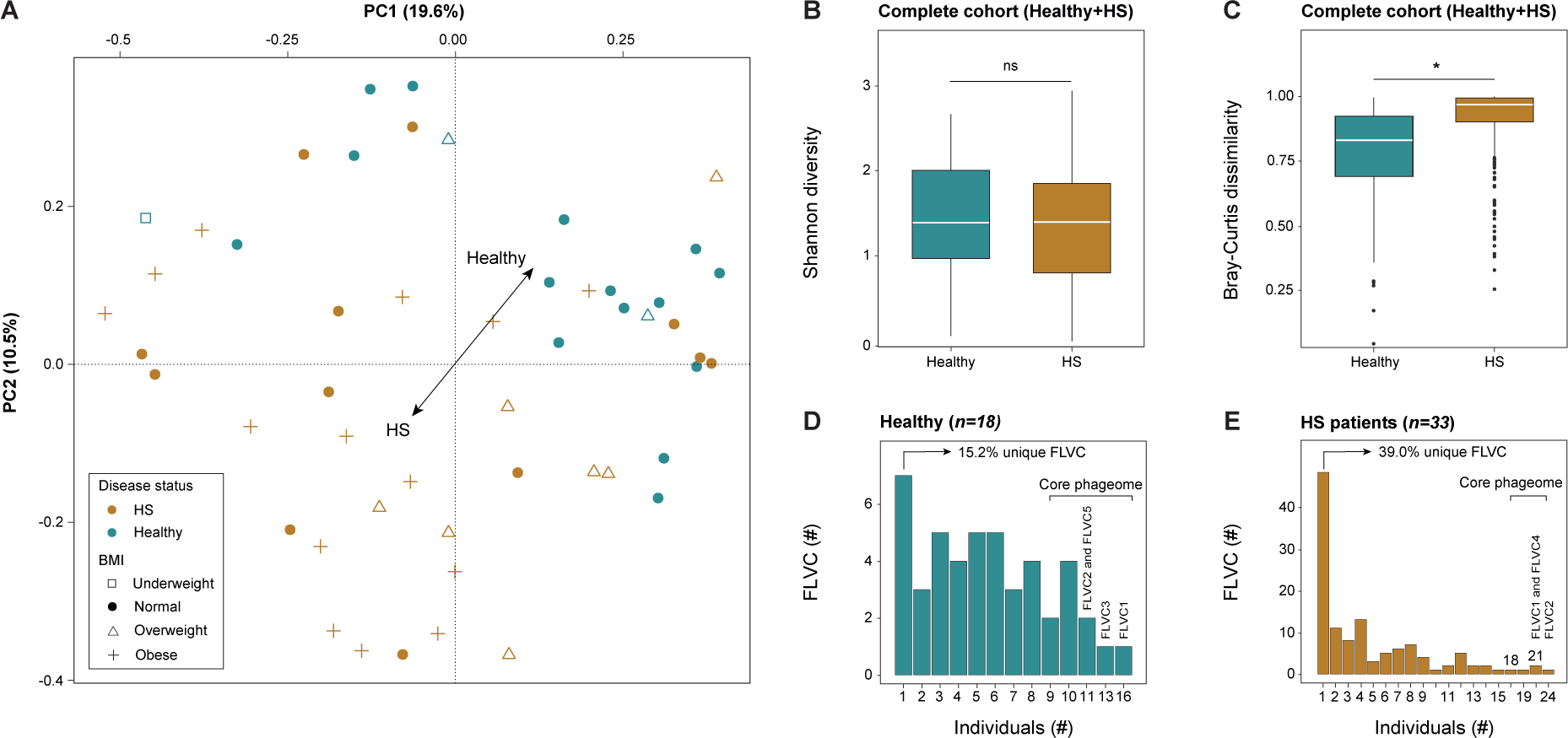
Skin virome diversity and viral sharing in the complete study cohort. **A,** A principal Coordinate Analysis visualizing the inter-individual differences of the skin virome composition (*n=51*, FLVC level, Bray-Curtis dissimilarity) of the complete cohort (Healthy+HS) coloured by disease status and with shapes depicted by categorical BMI. **B,** Alpha diversity (Shannon index) boxplot stratified according to disease status in complete cohort (*n=51*, FLVC level, Mann-Whitney U, AdjP≥0.05). **C,** Beta diversity (Bray-Curtis dissimilarity) boxplot stratified according to disease status in complete cohort (*n=51*, FLVC level, BMI=normal, Mann-Whitney U, AdjP< 2.2e-16). **D,** Barplot showing the absolute and relative number of FLVC shared between healthy individuals (*n=18*). **E,** Barplot showing the absolute and relative number of FLVC shared within HS patients (*n=33*). The core virome is defined by FLVC shared between 50% or more of healthy individuals or HS patients (most shared viruses are listed). Multiple testing adjustment (Benjamini-Hochberg method) was performed and significant associations (AdjP<0.05) are represented by an asterisk (*). Abbreviations: Hidradenititis suppurativa (HS)/Healthy controls (Healthy), Complete cohort (Healthy+HS), viral cluster (FLVC) and non-significant (ns).

**Extended Figure 6:**
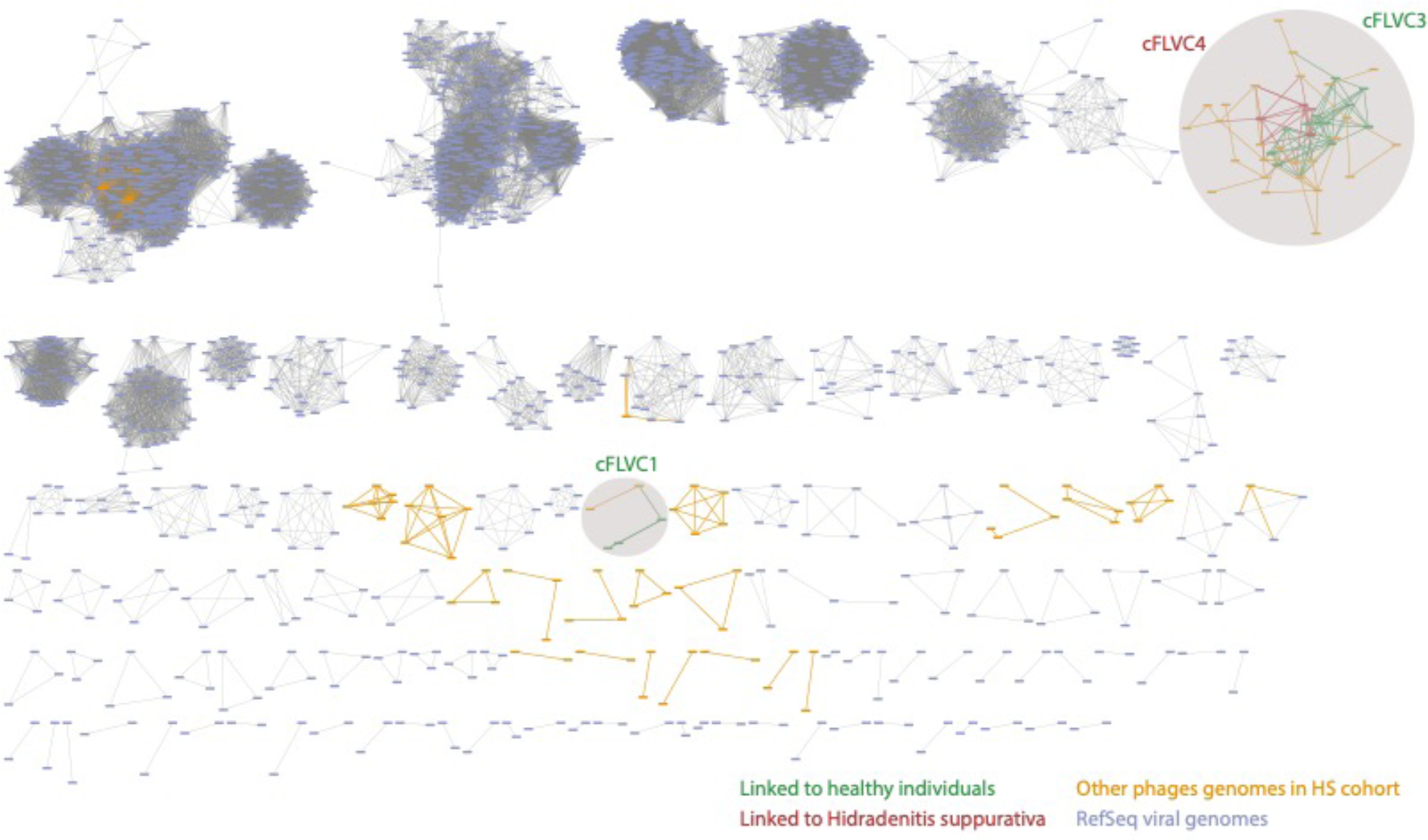
vConTACT2 clustering method for grouping phages at a genus-like level. Illustration of viral clusters found after combining the high-quality phages (*n=247*) with the current RefSeq viral database (*n=4406*). The nodes and edges in the network of the three viral clusters (FLVC1, FLVC3 and FLVC4) that are capable of differentiating between disease status and disease severity are colored (green/red) and highlighted with a gray circle. There was not a single connection found between the aforementioned viral clusters and the genomes present in the current RefSeq viral database (i.e., no single RefSeq genome within gray encircled area). There was also not a single connection found between the high-quality phages within FLVC3 and FLVC4 as they produced their own vConTACT2 clusters (Supplementary Table 12). Abbreviations: family like Viral cluster (FLVC).

**Extended Figure 7:**
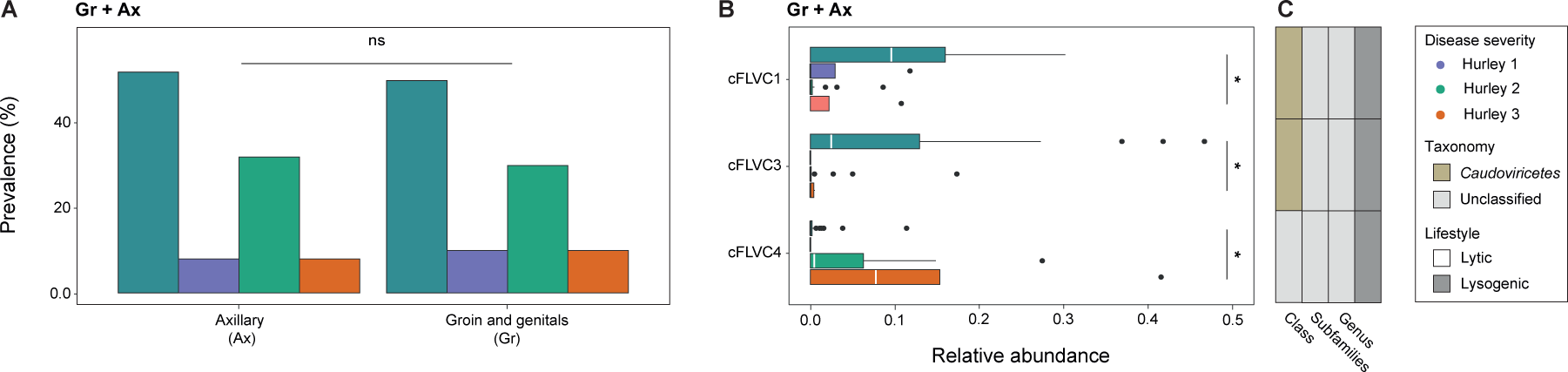
The relative abundance of the core virome in the location-deconfounded baseline cohort. **A,** The association between disease severity and location (axillary and groin and genitals) in the baseline sample cohort (*n=55*, chi-squared test, AdjP≥0.05). **B,** Relative abundance boxplot of the location-deconfounded core virome (present in axillary and groin and genital region) stratified according to disease severity in the baseline sample cohort (*n=55*, FLVC level, Kruskal-Wallis, AdjP<0.05). **C,** Classification (AS≥0.1) and lifestyle of representative high-quality phages of core viral clusters at class, subfamily and genus taxon. Multiple testing adjustment (Benjamini-Hochberg method) was performed and significant associations (AdjP<0.05) are represented by an asterisk (*). Abbreviations: Alignment score (AS), Hidradenititis suppurativa (HS)/Healthy controls (Healthy), complete cohort (Healthy+HS), axillary (Ax), groin and genitals (Gr) and non-significant (ns).

**Extended Figure 8:**
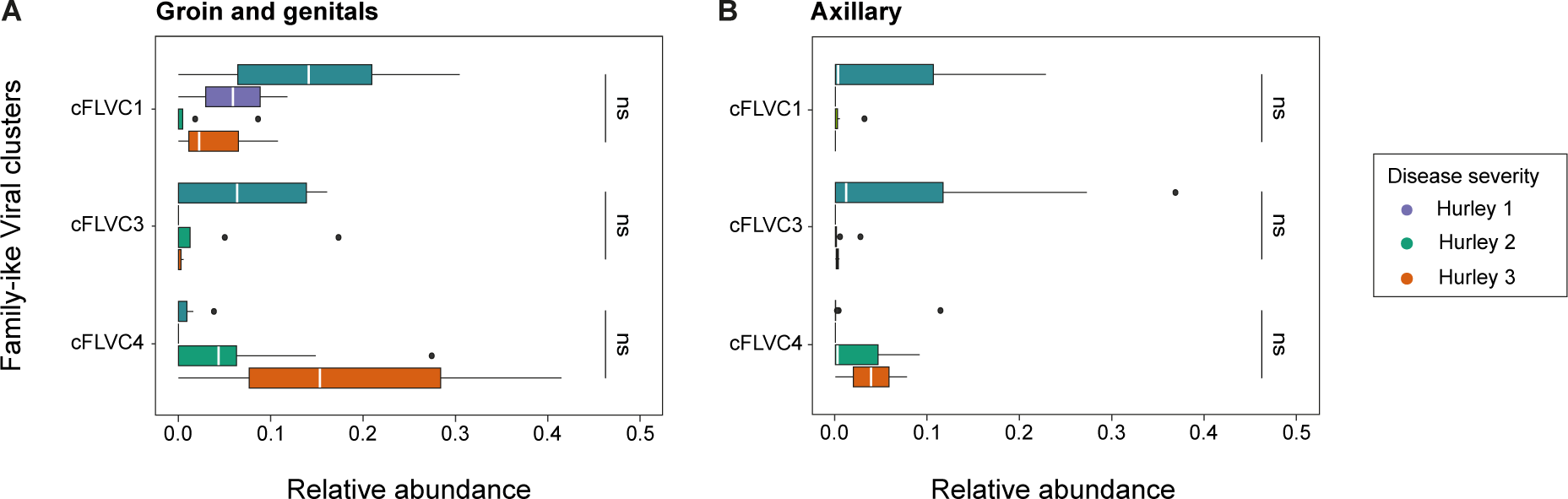
Relative abundance of core viral clusters affecting specific skin sites. **A,** Relative abundance boxplots of three core viral clusters stratified according to disease severity at groin and genital region in baseline sample cohort (*n=30*, Mann-Whitney U, AdjP≥0.05). **B,** Relative abundance boxplots of three core viral clusters stratified according to disease severity at the axillary region in baseline sample cohort (*n=25*, Mann-Whitney U, AdjP≥0.05). Adjustment for multiple testing was performed using the Benjamini-Hochberg method. Abbreviations: family like Viral cluster (FLVC) and not-significant (ns).

