## Extended Data Methods for "Hidradenitis Suppurativa Patients Exhibit a Distinctive and Highly Individualized Skin Virome"

#### Bioinformatic processing

The bioinformatic processing was carried out on a total of 4.1 billion paired end raw reads (equivalent to 0.62 TB), with some modifications from the previously described methodology (Extended Data Figure 1,2). The average number of raw reads per sample was 28.6 million. The raw reads were trimmed using Trimmomatic v0.39 to remove ambiguous bases, low-quality bases, and adapter sequences [1]. The resulting trimmed reads were then decontaminated by aligning them to both contaminome sequences constructed from negative controls and the reference human genome (hg38, BioProject=PRJNA31257) using bwa-mem2 v2.0 [2]. The average number of remaining quality-controlled reads per sample was 13.1 million. *De novo* assembly of the quality-controlled reads was performed using MetaSPAdes v3.15.1 with default parameters, resulting in a set of long contiguous sequences, known as contigs [3]. To mitigate the fragmentation of contigs with high coverage, a triple assembly approach was implemented by repeating the assembly step two additional times with a subset of the reads, namely 10% and 1% of the total reads. After the completion of the three assemblies, the contigs from each sample were merged and subjected to a first round of clustering (95% ANI and 85% coverage) using CheckV's scripts to remove redundancy [4]. Contigs shorter than 1,000 bp were discarded. A second round of clustering (95% ANI and 85% coverage) was conducted across all samples to further remove redundancy and produce a final non-redundant (NR) contig set (*n=52,429*), consisting of the largest contig present in each cluster. To calculate the abundances of each sample with high precision, we mapped quality-controlled reads to the NR contigs using bwa-mem2 v2.0, but only when the corresponding sample contained a member within the cluster of contigs represented by NR contigs [2]. To further prevent false positive phage identification and correct for spurious mapping, the abundance of each phage contig in a given sample was set to zero, if their horizontal coverage was below 70% [5]. By applying all the above mentioned quality checks, 17 samples (out of 144) were excluded, leaving a total of 127 samples for further analysis. The average number of mapped quality-controlled reads per sample was 3.06 million.

**Eukaryotic viruses**

The identification and classification of eukaryotic viruses was based on various homology-based approaches that relied on well-annotated public databases and can be found in Extended Data Figure 4. The first step involved comparing the NR-contig set against the NR protein database (August 29, 2022) using DIAMOND (sensitivity mode) v2.0.11 and CAT v4.6 [6,7]. The second step was to compare the NR-contig set against the NCBI nucleotide database (August 29, 2022) using BLASTN v2.11.0 (e-value ≤ 1e-10) [8]. The last step involved the final classification based on the principle of the lowest common ancestor using the ktClassifyBLAST module in KronaTools v2.7.1 [9]. To eliminate false positives, any classification with an alignment score (AAI/ANI x query coverage) below 0.1 were deemed unclassified.

**Prokaryotic viruses**

The identification of prokaryotic viruses (dsDNA, ssDNA or RNA), specifically bacteriophages or phages, was conducted using VirSorter2 v2.2.3 (--min-score ≥ 0.5) [10]. CheckV v0.8.1 was used to estimate the completeness of bacteriophage genomes [4]. Bacteriophages identified with Virsorter2 and an adequate quality tier (dsDNA, ≥ 50% completeness) and minimum genome length (ssDNA/RNA, ≥ 3kb) were labeled as “high-quality phages” and selected for further analysis. Next, a suite of homology-based and marker gene approaches was used to classify the selected bacteriophages [6–8]. Homology-based approaches previously described for eukaryotic virus classification were applied to bacteriophages as well. Phage classification was expanded by implementing a marker gene approach using Cenote-Taker2 v2.1.2 [11]. In addition, the bacteriophages were compared with a custom nucleotide database using BLASTN (e-value ≤ 1e-5, %cov ≥ 10,000 bp) to classify *Crassvirales*, as described earlier [5]. To confirm the novel character of the identified bacteriophages (not classified with aforementioned methodologies), we run vConTACT2 [12] together with the RefSeq database (April 14, 2023). Bacteriophages that did not cluster with a single RefSeq reference in vConTACT2 were considered 'newly described' high-quality phages, whereas those that clustered with a single reference were considered 'previously described' high-quality phages.

The lifecycle of bacteriophages was determined based on the presence of lysogeny-specific genes, which were predicted using the functional annotation module of Cenote-Taker2 (Supplementary Table 2) [11]. This results in the identification of genes necessary for a lysogenic life style necessary in 34.8% of the phage contigs. This functional annotation module uses Prodigal [13] and PHANOTATE [14] to extract open reading frames, which are then annotated by comparing to multiple of databases (CDD, Pfam, custom viral HMMs and PDB). The *in silico* prediction of bacterial host phyla (--min_cutoff ≥ 0.14) and genera (--min_cutoff ≥ 0.40) was performed using RaFAH v0.3 [15]. Additionally, the *in silico* predicted bacterial hosts were compared with the microbial 16S abundance profiles on genus level, described in another paper [16].

Although the viral identification approach described above was capable of capturing high-quality phages, they only represented 40.2% of all viral reads (Extended Data Figure 2). This was likely due to the fragmented and/or diverse nature of skin phages that are currently underrepresented in viral databases, as suggested before [17]. To increase the fraction of viral reads in our further analyses, we implemented the following viral cluster (VC) approach (allowing the inclusion of reads mapping to phage contigs which were not identified as 'high quality' using CheckV):

1. Exclude NR-contigs that were identified as a eukaryotic virus.
2. Exclude NR-contigs shorter than 3,000 bp.
3. Cluster NR-contigs based on a combination of gene sharing and pairwise average amino acid identity, as described by *Nayfach* and colleagues [18], resulting in "family-like viral clusters" (FLVC).
4. Mark all contigs present in FLVC containing at least one high-quality phage, along with their representing NR-contigs, as viral.

This FLVC approach increased the percentage of viral reads included in our analyses to 66.9% (Extended Data Figure 2), encompassing a total of 126 phage family-like clusters (mean number of clusters present per sample=11), indicating a comprehensive representation of the skin virome. Additionally, the FLVC classification was performed based on the principle of the lowest common ancestor of all the high-quality phages present within a given FLVC (Supplementary Table 2). Finally, 18 samples were discarded due to the absence of high quality phage genomes, leaving a total of 109 samples for further viral analysis. The average number of viral reads per sample was 2.70 million.

**Skin virome features and visualization**

Virome intra-individual diversity was calculated by various alpha-diversity metrics

(Shannon diversity and observed richness) on the family abundance table using the phyloseq R package [19]. Virome inter-individual variation, better known as beta-diversity, was determined using Bray-Curtis dissimilarity on the family abundance table with Hellinger transformation and visualized by principal coordinate analysis (PCoA). Considering the sparse nature of virome data and the resulting significant data loss, we opted not to use CLR-transformation (default 10% prevalence threshold) followed by principal component analysis visualization.

**Skin virome compositional variation across clinical metadata variables**

Univariate distance-based redundancy analysis (dbRDA) was performed using the capscale function from the vegan R package to determine the effect of metadata on the virome variation (family level, Bray-Curtis) [20]. The metadata variables that demonstrated a significant contribution to the virome variation in the univariate analysis were exclusively considered for following multivariate analysis. Multivariate dbRDA with forward model selection was performed using the *ordiR2step* function in vegan to determine the non-redundant cumulative effect of the metadata on the virome variation (family level, Bray-Curtis). The contribution of significant metadata variables on the first two principal coordinates were determined using the *envfit* function, as implemented in the vegan package (univariate dbRDA) and were plotted as arrows on the PCoA plot [20].

**Statistical analysis**

Statistical analysis was performed in R using the stats, vegan and phyloseq packages [19–21]. The statistical tests were two-sided, non-parametric, with a significance level set at $P < 0.05$. Appropriate multiple testing correction was carried out using the Benjamini-Hochberg (BH) method, where significance was defined as $AdjP < 0.05$. The rstatix package was used to compute the Wilcoxon effect size $r = Z/\sqrt{N}$ and Chi-squared effect size $r = \sqrt{\chi^2}/N$.

### Code availability

The Virome Paired-End Reads (ViPER) pipeline was used for the bioinformatic processing of the raw paired end reads (https://github.com/Matthijnssenslab/ViPER). For the purpose of reproducibility, additional data can be accessed at https://github.com/Matthijnssenslab/IBDVirome/tree/main/IBDHS.

### Data availability

Supplementary Table 1 contains the clinical metadata. The raw sequence data have been deposited to the NCBI Sequence Read Archive with BioProject accession number PRJNA961962. The high-quality phage genomes of cFLVC1, cFLVC3 and cFLVC4 been deposited to GenBank with the accession numbers OQ890309-OQ890326, but due to a high submission volume, they are still being processed and are accessible via Figshare: https://figshare.com/projects/The_skin_virome_in_Hidrandenitis_suppurativa/165766

21. R Development Core Team R. R: A Language and Environment for Statistical Computing. R Foundation for Statistical Computing. 2011.
